# Tracking the relation between gist and item memory over the course of long-term memory consolidation

**DOI:** 10.1101/2021.01.05.425378

**Authors:** Tima Zeng, Alexa Tompary, Anna C. Schapiro, Sharon L. Thompson-Schill

## Abstract

Our experiences in the world support memories not only of specific episodes but also of the generalities (the ‘gist’) across related experiences. It remains unclear how these two types of memories evolve and influence one another over time. 173 human participants encoded spatial locations from a distribution and reported both item memory (specific locations) and gist memory (center for the locations) across one to two months. After one month, gist memory was preserved relative to item memory, despite a persistent positive correlation between them. Critically, item memories were biased towards the gist over time; however, with a spatial outlier item, the local center excluding the outlier became the source of bias, instead of the reported center overweighting the outlier. Our results suggest that the extraction of gist is sensitive to the regularities of items, and that the gist starts to guide item memories over longer durations as their relative strengths change.

## Introduction

Our experiences in the world are perceived and remembered both as individual items, events, and episodes, and also as aggregated collections or sets of related items with common properties. A fundamental question in cognitive science is how we extract summary statistics from individual instances, both during perception and working memory (where aggregated information is often referred to as an “ensemble”) and during episodic encoding and retrieval (where aggregated information is often referred to as a “schema” or as the “gist” of an experience). In addition, researchers have attempted to characterize how memory for these types of information changes over time. For example, studies of long-term memory in both humans and animals have demonstrated that gist memory persists or even improves over time, whereas memory for the individual items from which the gist is built fades (Posner & Keele, 1970; Richards et al., 2014).

What do these observations of temporal dissociations tell us about the relation between item memory and gist memory? On the one hand, a persisting gist memory with less accurate item memory is often taken as evidence that a gist representation becomes independent of individual item representations as it is abstracted during encoding (Posner & Keele, 1970) or through consolidation (Richards et al., 2014). On the other hand, a persisting gist memory with less accurate item memory is not sufficient evidence for the independence of a gist representation: Even when item memories become noisy and less accurate, they still can retain enough information to support a relatively intact memory of gist at retrieval (Alvarez, 2011; Squire, Genzel, Wixted, & Morris, 2015).

Disentangling these two possibilities based on existing evidence is difficult, because previous studies do not have a direct measurement of the gist information retained in item memories. In this study, we developed a paradigm to test item memory, gist memory, and “estimated” gist memory, which is an estimate of gist memory given the assumption that it is assembled from individual memories of constituent items. Ensemble perception research was a source of inspiration in developing such a paradigm. Studies of rapid perception of complex visual arrays reveal precise representations of gist (i.e., ensemble statistics) with less accurate item memory retrieval in working memory (Ariely, 2001). In order to investigate the relation between item and gist, ensemble perception paradigms often operationalize the gist as the average representation across instances.

In our paradigm, participants learn a set of landmarks (e.g. restaurant) whose locations are clustered together, and then they are tested on the locations of each landmark (item memory) as well as their spatial center (reported gist). Importantly, an “estimated center” can be computed based on their retrieval of individual items, and its accuracy can thus reveal the amount of gist information available in item memories. Thus, we can investigate the relation between gist memories and item memories over the course of long-term memory consolidation by measuring the relationship between the accuracy of the reported center and the estimated center. A positive correlation between estimated and reported center accuracy could mean that participants’ gist memory was still supported by individual item memories, or that the gist was influencing the retrieval of items.

To probe the direction of this relationship, we developed a gist-based bias measurement, an approach borrowed from research on hierarchical clustering models, that reveals how much gist memory influences memory for specific items (Brady & Alvarez, 2011). Theories suggest that this influence reveals a reconstructive memory retrieval process (Brady, Schacter, & Alvarez, 2015; Hemmer & Steyvers, 2009; Schacter, Guerin, & Jacques, 2011) that depends on the relative strength of item and gist memory (Tompary, Zhou, & Davachi, 2020). Consistent with this theory, prior work in long-term memory consolidation has shown that over time, as the strength of gist memory is preserved or improves and/or that of item memory decreases, items that are consistent with the gist are recalled more precisely (Richter, Bays, Jeyarathnarajah, & Simons, 2019;Sweegers & Talamini, 2014; Tompary, Zhou, & Davachi, 2020). However, these results are different from a gist-based bias, a distortion of item memory. An increasing bias of item memories towards the remembered gist — in this paradigm, the reported center — would be strong evidence for the increasing strength of gist memory over item memories. Our interest in examining the influence of gist memory on item memory at long delays stems from a desire to bridge the literature reviewed above with a potentially related literature reporting the effects of prior knowledge on memory retrieval (e.g., Huttenlocher, Hedges, & Vevea, 2000; Tompary & Thompson-Schill, 2019).

The current study aimed to understand the relation between item and gist memory over the course of a month. In Experiment 1, we trained three groups of participants on spatial locations of six landmarks, and we measured the change in memory of these items as well as the memory for the gist (i.e. the center participants reported) at one of three delay periods: 24 hours, one week, or one month. We predicted that the accuracy of the reported center would persist or improve despite the accuracy of retrieved items decreasing over a month, as seen in prior work. We extended prior observations by including two new measures—estimated center and gist-based bias—in order to explore how the relation between memories for items and the gist change over the course of one month.

In Experiment 2, we explored the influence of an “outlier” item in spatial location both on the gist, and on the relation between item and gist memories demonstrated in Experiment 1. Research in long-term memory (Richards et al., 2014) and working memory (Whitney & Leib, 2018) shows that outliers that are inconsistent with the pattern across all items differently influence the memory for the pattern, compared to items that are more consistent with the pattern. Outliers greatly disrupted or shifted the overall pattern (Richards et al., 2014), or were discounted in estimating the pattern (Haberman & Whitney, 2010) compared to other items. In Experiment 2 we examined the extent to which the gist representation was influenced more or less by an outlier item over time, as well as whether bias in item memory was to the center location including or excluding the outlier item. Taken together, these experiments provide new information about how item and gist memory are consolidated over time.

## Results

**Experiment 1** Item memories were operationalized as “landmarks” (i.e. dots associated with unique landmark names) in six locations on a laptop screen (Fig. 1a). In Session 1, 130 participants learned the locations of the landmarks individually through a training to criterion procedure (see Fig. 1b and Methods for details). After training, participants were tested on item and gist memory: First, they indicated their guess about the center of the landmarks (gist memory test), and then they recalled each landmark location, without feedback and in a random order (item memory test). After 24 hours (*n* = 44), one week (*n* = 43), or one month (*n* = 43), participants returned for Session 2, during which they completed the gist memory test followed by the item memory test again. This testing order was chosen to reduce the influence of item memories on reported gist.

**Fig. 1.**
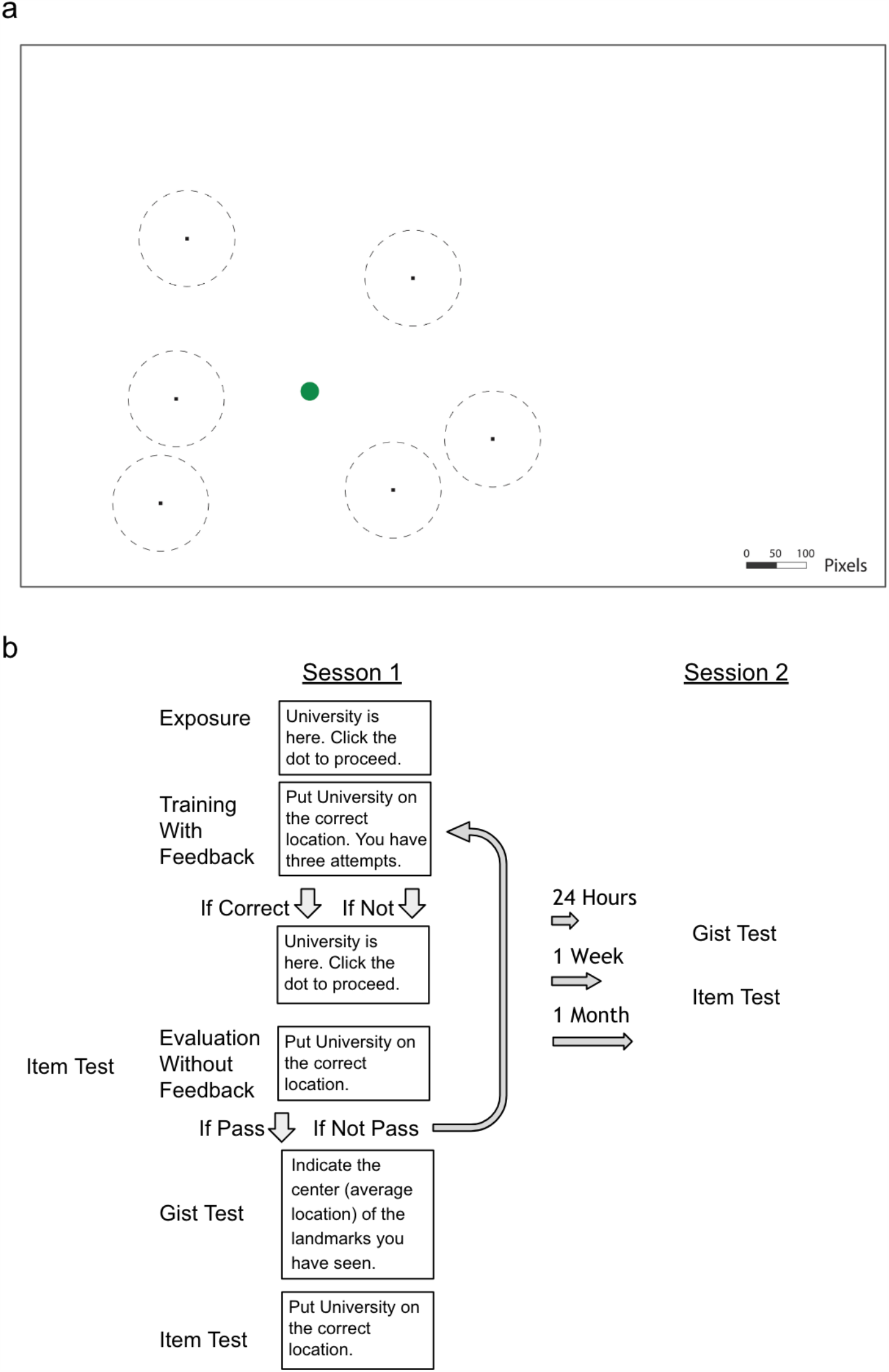
**a)** Illustration of item locations, drawn to scale. The locations (black dots) were the same for all participants, but the mapping between the location and landmark name was randomized for each participant. The dash lines around the dots indicate the training criteria (80 pixels). The green circle indicates the center of these encoded locations. **b)** Procedure. Participants completed cycles of encoding (with feedback) and evaluation (without feedback) until they could retrieve each landmark individually within the training criteria.

### Gist memory decreased less than item memory over time

As described above, we developed an error measurement for the accuracy of item and gist memory (Fig. 2a; See Methods for details). All three delay groups performed above chance on both the item memory test (chance defined as the average of distance between encoded item locations and center of the screen) and the gist memory test (chance defined as the distance between the center of encoded locations and center of the screen) at both Session 1 and 2 (all p< .0001; Supplementary Fig. 1). To examine how item and gist memory changed over time,, we conducted a 3 (group: 24-hour, 1-week, and 1-month) X 2 (memory type: item, center) aligned ranks transformation ANOVA of the difference in error between Session 1 and Session 2 (because the data were not normally distributed). This test revealed a main effect of group, *F*(2, 254) = 42.26, *p* < .001, a main effect of memory type, *F*(1, 254) = 99.36, *p* < .001, and an interaction between group and memory type, *F*(2, 254) = 23.76, p < .001. This interaction reveals that the error in retrieved items increased more over time compared to error in the reported center (Fig. 2b). Specifically, whereas each pairwise comparison between groups was significant for item memory (Mann-Whitney tests: all *p*’s < .01), the only reliable group difference for gist memory was between the 24-hour and 1-month groups (*U* = 685, *p* = .026). In addition, the change in retrieval error of item locations was significantly higher than that of reported center at a delay of 24 hours (Wilcoxon signed rank tests: *Z* = 2.14, *p* = .03), one week (*Z* = 3.06, *p* < .01), and one month (*Z* = 5.32, *p* < .0001). These results showed that item memories decreased significantly more relative to gist memory over time.

**Fig. 2.**
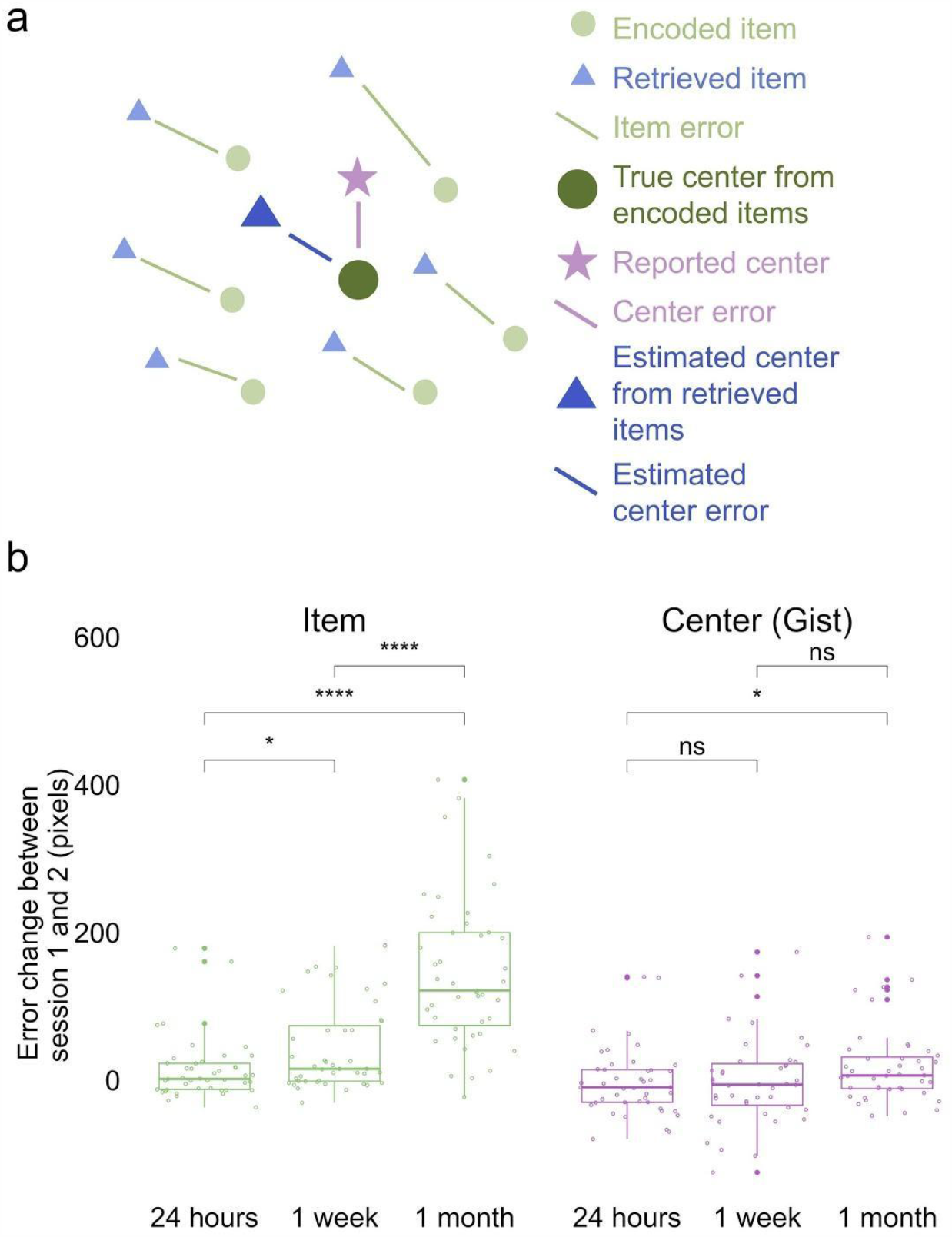
Error measurements and results. **a)** Error measurements. **b)** Change in error by group and memory type (the band indicates the median, the box indicates the first and third quartiles, the whiskers indicate ± 1.5 × interquartile range, and the solid points indicate outliers). Greater values indicate an increase in error in Session 2 over Session 1. * indicates *p* < .05 and **** *p*< .0001.

### Positive relationship between item and gist memories across one month

To explore the relation between item and gist memory over time, we used a linear model to evaluate the effects of estimated center error, delay group, and their interaction on reported center error. We found that estimated center error significantly predicted reported center error (*b* = 0.73, *t*(69.92) = 3.48, *p* < .001). The effect of delay (*b* = −10.45, *t*(59.59) = −0.77, *p* = .45) and the interaction between the estimated center error and delay (*b* = −0.20, *t*(68.80) = −0.68, *p* = .50) on reported center error was not significant. These results indicate a stable relation between item and gist memory over time; our subsequent analyses will examine the source of this relation.

### Item memory retrieval biased towards gist over time

The positive correlation could indicate that participants’ reported gist was still supported by individual item memories, or alternatively that the reported gist was influencing the retrieval of items. To examine the direction of the relation between item and gist memories, we developed a bias analysis (controlling for the magnitude of error; see Methods for details) as a measure of how much the retrieval of each item memory is biased towards the gist representation (Fig. 3b). We used the reported center (not the true center) in this analysis because we found a decrease in accuracy of the reported center after one month compared to 24 hours, as discussed above^1^.

**Fig. 3.**
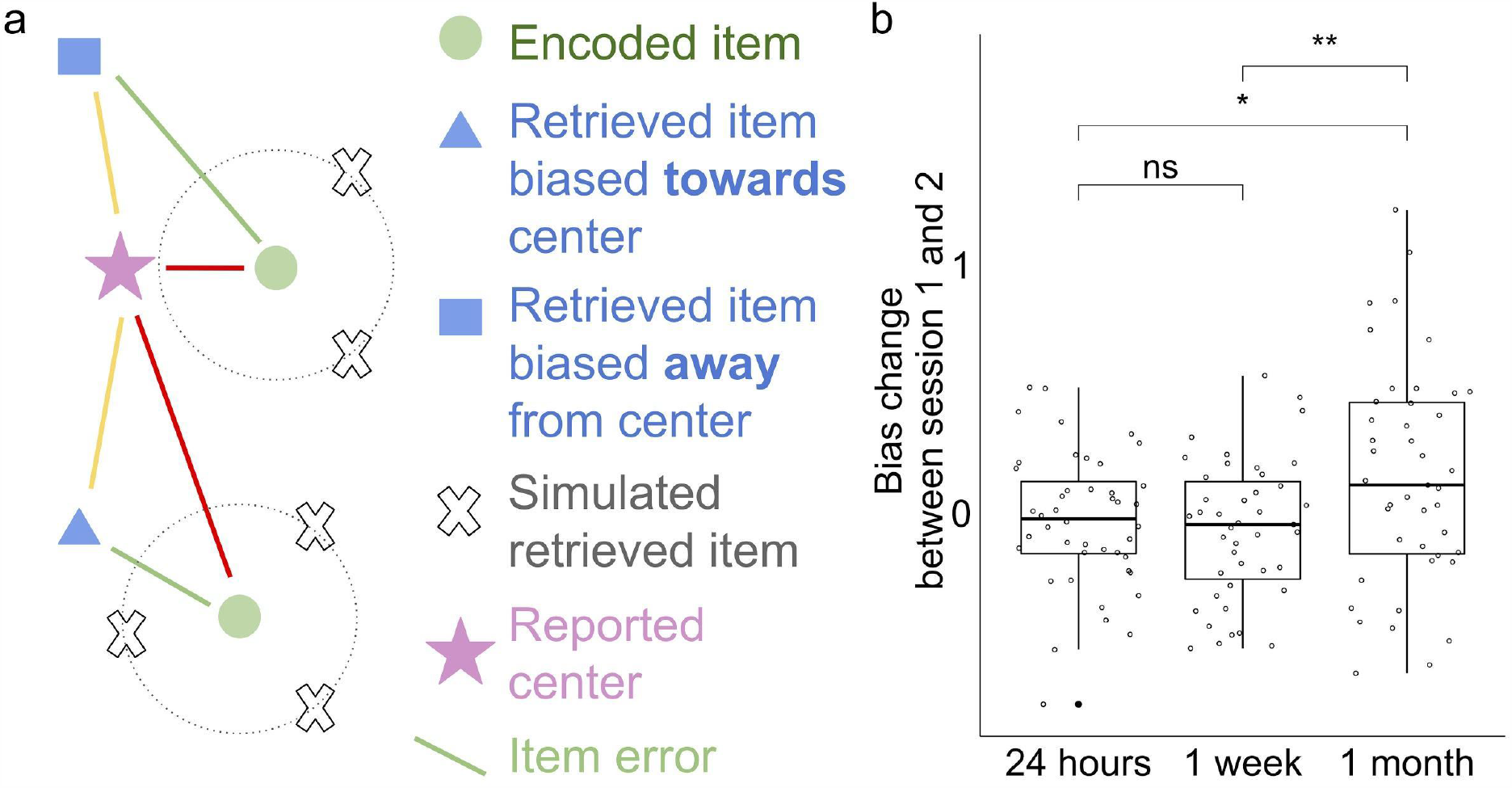
**a)** Bias measurements. The absolute bias for each location is (red - yellow) / green.The blue square is an example of a recalled item that is biased away from the reported center and the blue triangle is an example of a recalled item that is biased towards the reported center. Bias for each participant is an average of absolute bias for all the locations minus baseline bias (i.e. a bias value calculated from simulations of item memories with the same errors as the data but assuming no directions). **b)** Change in recalled items’ bias towards the reported center for the 24-hour, 1-week, and 1-month groups (the band indicates the median, the box indicates the first and third quartiles, the whiskers indicate ± 1.5 × interquartile range, and the solid points indicate outliers). Values > 0 indicate item memory was more biased towards the reported center in Session 2 relative to Session 1. * *p* < 0.05 and ** *p* < 0.01 by t-tests.

Only the bias value at 1 month was significantly greater than 0, *t*(42) = 3.69, *p* < .001. We conducted a one-way ANOVA of change in bias (from Session 1 to Session 2) across the three groups to examine if participants’ item memory became more biased towards the reported center over time. This test revealed a main effect of group, *F*(2, 127) = 5.00, *p* < .01. Pairwise between groups comparisons revealed a greater change in bias between the 24-hour and 1-month groups, *t*(68.74) = 2.29, *p* = .02, and also between the 1-week and 1-month groups *t*(71.13) = 2.69, *p* < .01, but not between the 24-hour and 1-week groups, *t*(84.54) = 0.59, *p* = .55. These results indicate that by one month (but not after one day or one week), item memory retrieval was biased towards the remembered gist.

Taken together, the results of Experiment 1 reveal that although there is a persistent relation between item and gist memory during memory consolidation, the nature of this relation changes over time. We suggest that early in memory consolidation, retrieval of gist depends on the successful retrieval of individual items, but then, as item memory weakens over time, the relatively stronger gist memory begins to guide retrieval of item memory. As a consequence, this new gist representation can exert influence over memories in ways described by reconstructive memory theories.

**Experiment 2** The stimuli and procedure of Experiment 2 (see Fig. 4) were similar to those of Experiment 1 but differed in two major ways (see Methods for more details). Firstly, we used a repeated measures design in Experiment 2 so that we could observe changes in memory at short (one day) and long (1-2 month) retention intervals within each subject (N = 43). Secondly, one of the landmarks was an “outlier”, meaning that its location fell far out of the range of the cluster where the majority of the landmarks were (e.g., “Park” in Fig. 4a). The inclusion of an outlier location enabled us to examine the influence that a single “atypical” item has not only on the initial estimation of the center (as in Haberman & Whitney, 2010; Richards et al., 2014; Whitney & Leib, 2018) but also on the source of bias in item memory at long delays. In addition, in Session 1, item memory was derived from the item memory test in the last round of evaluation during training (Fig. 4b) to streamline the session.

**Fig. 4.**
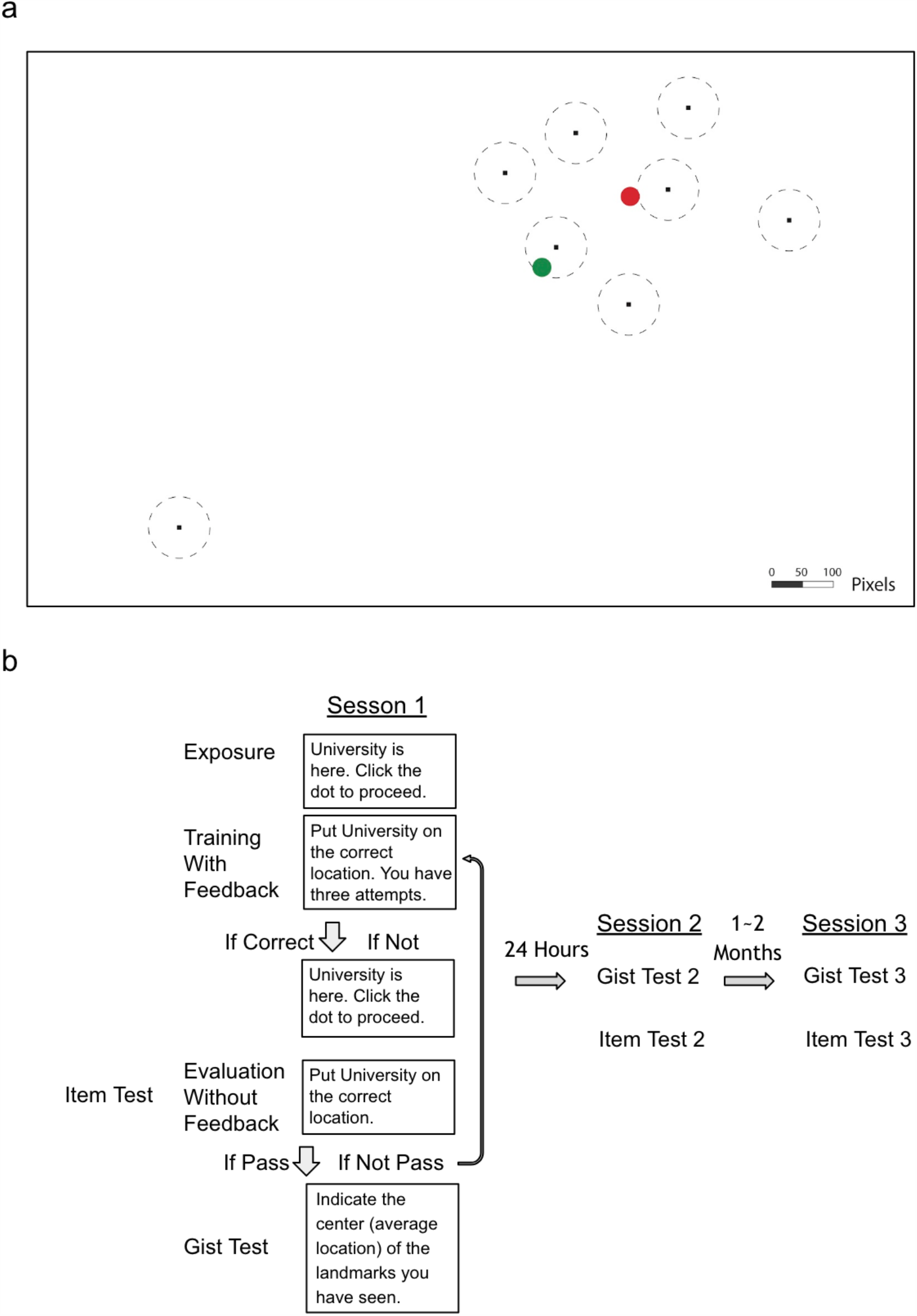
**a)** Experiment 2 stimuli drawn to scale. The green circle indicates the global center of all encoded items. The red circle indicates the local center of encoded items excluding the spatial outlier. Dots indicate the locations of landmarks. Dash lines around the dots indicate the training criteria (50 pixels). **b)** Experiment 2 procedure.

### Gist memory decreased less than item memory over time

In Experiment 2, we used the same error measurement for the accuracy of item and gist memory as in Experiment 1 (Fig. 2a; see Methods for details). Participants performed above chance in both item (i.e. the average of distance between encoded item locations and center of the screen) and gist memory (i.e. the distance between the center of encoded locations and center of the screen) tests at all sessions (all p < .0001). To examine how item and gist memory changed over time, we conducted a 2 (delay: short (24 hours) or long (1-2 months) x 2 (memory type: item, center) aligned ranks transformation ANOVA with repeated measures of error change. This test revealed a main effect of delay, *F*(1, 126) = 16.20, *p* < .001, memory type, *F*(1, 126) = 68.96, *p* < .001, and an interaction between delay and memory type, *F*(1, 126) = 27.40, *p* < .001. This interaction indicates that the error of individual item retrieval increased more over time compared to the reported center (Fig. 5a). Specifically, whereas item memory error change was higher after 24-hour compared to one to two months by Wilcoxon signed rank test (*Z* = 5.28, *p* < .0001), no such significant difference was detected for gist memory error change (*Z* = 0.72, *p* = .47). Experiment 2 thus replicated the finding observed in Experiment 1 that item memories decreased significantly more relative to gist memory over time.

**Fig. 5.**
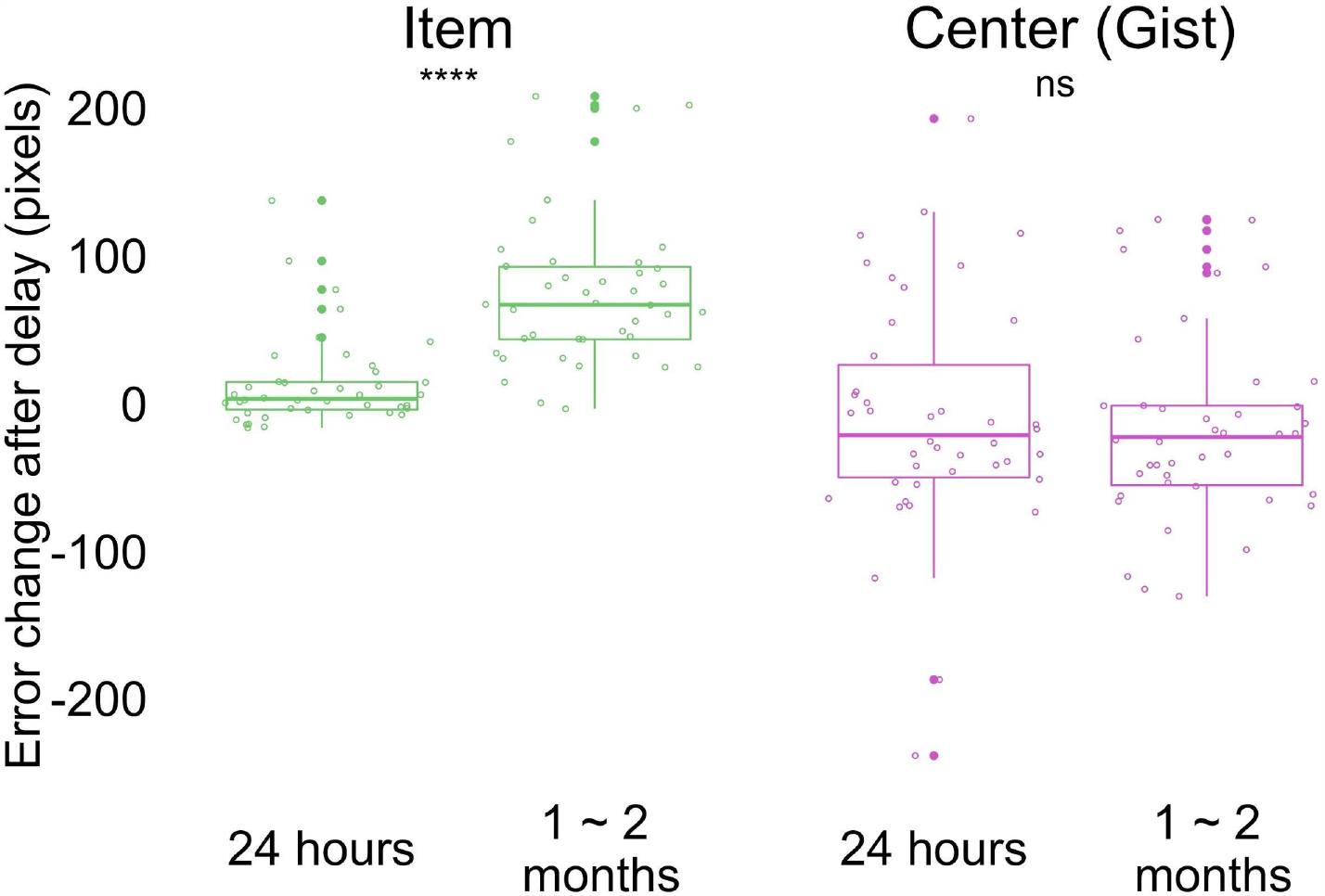
Change in error by delay and memory type (the band indicates the median, the box indicates the first and third quartiles, the whiskers indicate ± 1.5 × interquartile range, and the solid points indicate outliers). Greater values indicate an increase in error from Session 1 after delay. **** *p* < .0001.

### Positive relationship between item and gist memories across time

To explore the relation between item and gist memory, we fit a linear mixed effects model on reported center error with fixed effects of delay (24 hours and 1 to 2 months), estimated center error, and their interaction with a random effect of participant to account for repeated measures within participants. We found a significant effect of estimated center error (*b* = 0.73, *t*(69.92) = 3.48, *p* < .001), but not a main effect of delay (*b* = −10.45, *t*(59.59) = −0.77, *p* = .45) or an interaction between the estimated center error and delay (*b* = −0.20, *t*(68.80) = −0.68, *p* = .50). The result of a persistent relationship between estimated center error and reported center error at short and long retention intervals was replicated in a within-participants design.

### Item memory retrieval was biased towards the local gist over time

To examine whether the influence of gist on item memories changes over time, we applied the same bias measurement analysis controlling for errors as in Experiment 1, using participants’ reported center of all the retrieved items as bias center (Fig. 3b). In contrast to Experiment 1, the change in bias after a long retention interval was not significantly higher than 0 (*t*(42) = −0.43, *p* = .67). Bias after a long retention interval was marginally greater than bias after a short retention interval (*t*(42) = −1.70, *p* = .10) by paired t-test; however, this was driven by an unpredicted negative bias (i.e., bias away from the reported gist) immediately after learning (*t*(42) = −2.09, *p* = .04) and after a short retention interval (*t*(42) = −2.27, *p* = .03) by t-test against 0.

What might explain the different bias results between Experiments 1 and 2? We suspect this is the result of the outlier item. Prior work in visual working memory research showed that outliers were discounted in estimating the gist (Haberman & Whitney, 2010). In our Experiment 1, where there was not an outlier, we saw that the item retrieval was biased towards the center of all of the items; however, in Experiment 2, the center of *most* of the items would be the local center excluding the outlier. It is possible that for participants in Experiment 2, the items were biased towards the local clustering center excluding the outlier.

In order to test this possibility, we conducted an analysis that computed the gist-based bias of the items using the local center (i.e. the true center from the encoded items disregarding the outlier). As in Experiment 1, only the bias at the long retention interval was significantly greater than 0 (*t*(42) = 2.30, *p* = .03); furthermore, the bias change after a long retention interval was significantly higher than the bias change after a short retention interval (*t*(42) = 2.87, *p* < .01). This effect was observed even for the outlier item: Retrieval of the location of the outlier item was significantly more biased towards the local center after a long retention interval compared to a short retention interval (*Z* = 2.46, *p* = .01). These results indicate that item memories in Experiment 2 were biased, at long retention intervals, towards the center as in Experiment 1, but that the “center” in Experiment 2 was not the global center but instead the local center of the display.

### The outlier weighted more compared to other items in gist memory

Our analysis of item bias suggests that the outlier is “discarded” as a member of the cluster of locations, which is consistent with some prior studies (e.g., Haberman & Whitney, 2010); however, other work has shown that outliers can greatly disrupt or shift the representation of a set of events (Richards et al., 2014). Could both be happening in this paradigm? In order to explore the influence of the outlier on the representation of gist, we applied a weighted model adapted from working memory literature and computed an estimation of the weight of the outlier in the reported gist (Haberman & Whitney, 2010; see Methods for details).

The estimated weight of the outlier was significantly higher than 0.125 (i.e. the level assuming equal weights across all items) immediately after learning (t(42) = 2.14, p = .04), after a short retention interval (t(42) = 2.89, p < .01) and after a long retention interval (t(42) = 3.83, p < .001).The change in outlier weight after short compared to long retention intervals did not significantly differ (t(42) = 0.92, p = .36). In other words, the outlier has not been discarded from the set, but quite to the contrary, the outlier has a disproportionate influence on the explicit retrieval of gist after a delay. In contrast, the implicit effect of the center on bias in item retrieval seems to emerge from a center that is uninfluenced by the outlier (Fig. 6b). These results revealed that the outlier consistently influenced participants’ reported center more than other items at all tested time points.

**Fig. 6.**
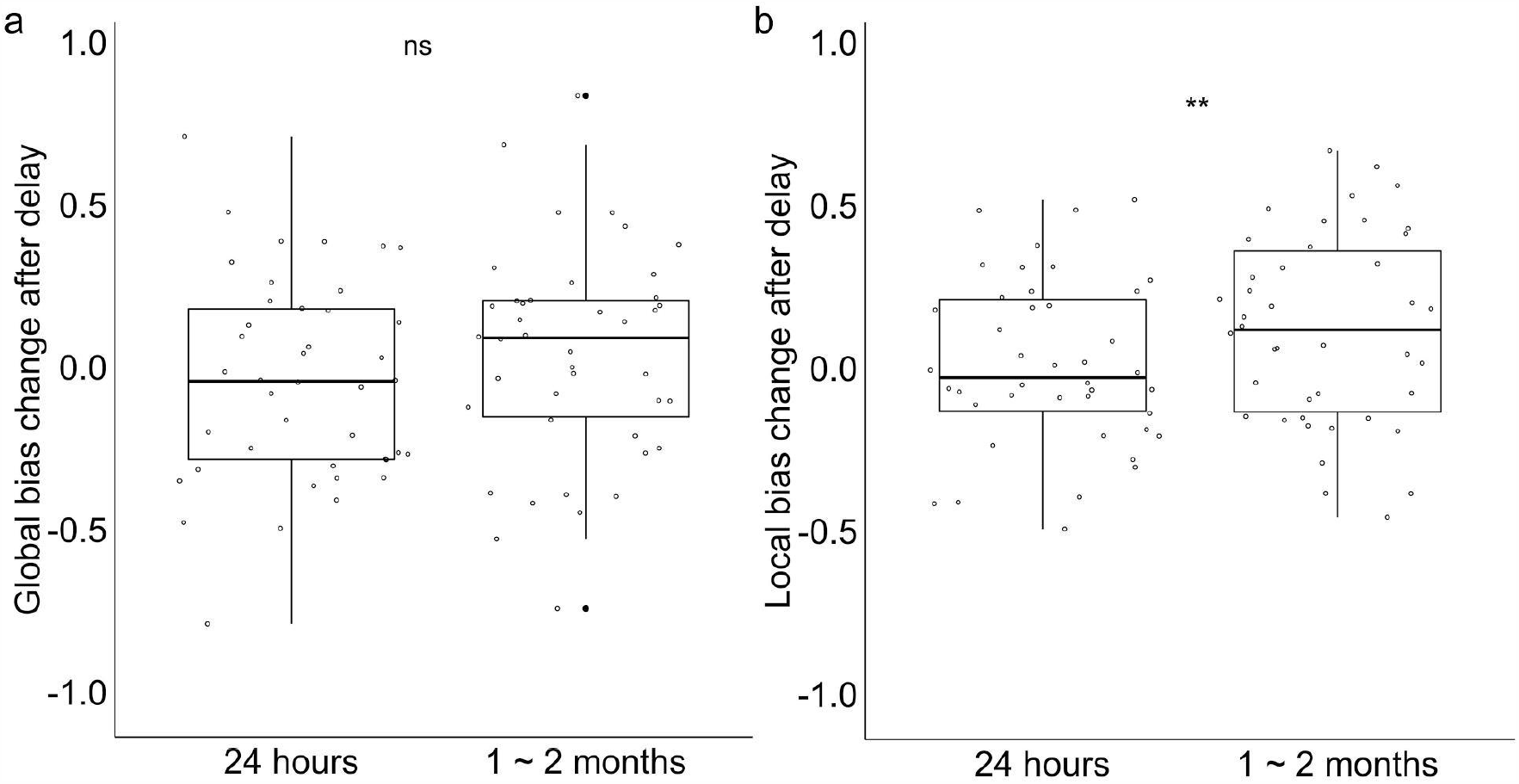
Change in item bias. **a)** Change in item bias towards the reported center after short (1 day) and long (1-2 months) retention intervals (the band indicates the median, the box indicates the first and third quartiles, the whiskers indicate ± 1.5 × interquartile range, and the solid points indicate outliers). **b)** Change in item bias towards the local center after short and long retention intervals. Values > 0 indicate item memory was more biased towards the local center relative to Session 1 after delay. ** *p* < 0.01.

Taken together, Experiment 2 replicated the main findings from Experiment 1 that gist memory decreased less compared to item memories over time (Fig. 5a) and a positive relationship between item and gist memories (Fig. 5b) in a within-subject design. The “outlier” item changed the relationship between items and reported global center after a long retention interval. By one to two months, items were no longer biased towards the global reported center which overweighted the outlier. Instead, they were biased towards the local center excluding the outlier over time.

## Discussion

We examined how human learners extract the “gist” (generalities, common properties, or summary statistics) across individual instances, and how memory for these instances and for the gist evolve and influence each other over time through two behavioral experiments spanning one to two months. We demonstrated that the accuracy of item memory (memory for spatial locations on a screen) decreased more compared to the accuracy of gist memory (center of the locations) over time, though there was a persistent positive correlation between these them. Critically, item memories were increasingly biased towards the gist over time. In the presence of an outlier item, the local gist, excluding the outlier, became the source of bias, instead of the gist participants directly reported, which consistently overweighted the outlier across time. We think that gist memories, initially built from item memory, gradually developed to guide item memory as their relative strength changed over time.

Consistent with prior research (Lutz, Diekelmann, Hinse-Stern, Born, & Rauss, 2017; Posner & Keele, 1970), item memory became less accurate over time while gist memory remained relatively intact over time. Our findings converge with this prior research even when using an explicit instruction to retrieve gist memory, rather than inferring gist from another measure as in prior research. A shortcoming of prior research is that the relation between item and gist memory over time is rarely assessed. We showed that the relationship between item memory and gist memory persisted across delay periods despite decreased accuracy in item memory. This relationship could have resulted from the influence of gist memory on the retrieval of item memory, from the influence of item memory on gist memory, or both. Our gist-based bias results shed light on the direction of this relationship: Item memory retrieval was biased towards gist only after one month, which suggested that the correlation at one month was likely to be due to the influence of gist memory on the retrieval of item memory.

Our findings that items are increasingly biased towards the gist as the accuracy of item memories decreases over time provide new evidence for the memory reconstruction framework, which proposes that memory retrieval is a combination of different sources with varying strength (Brady et al., 2015; Hemmer & Steyvers, 2009; Huttenlocher et al., 2000; Tompary et al., 2020). Our work extends prior evidence of increased schematization in memory consolidation (Richards et al., 2014; Richter et al., 2019; Tompary et al., 2020) by demonstrating a new form of influence from the gist on item memory: gist-based bias. In contrast to prior memory consolidation research that showed increased schematization earlier than one month (Richter et al., 2019; Tompary et al., 2020), the gist-based bias in our current work did not appear after 24 hours or one week. This discrepancy could be because the intensive training participants experiences in our paradigm increased the strength of item memories relative to gist memory during learning, and only after a long retention interval did the strength of item memory decrease to an extent that allowed bias to manifest. The results are also consistent with a slow systems consolidation process that results in a qualitatively different representation of these memories only after a longer retention interval of one month. These two types of memories interact dynamically (Richards et al., 2014; Winocur & Moscovitch, 2011; Sekeres, Winocur, & Moscovitch, 2018). An increased reliance on neocortical areas over time would be expected to strengthen gist memory, as neocortex tends to represent information in a ‘semanticized’ form. The pull of item memories toward the gist at one month may thus reflect the slow establishment or stabilization of such a neocortical trace.

Our results of increasing gist-based bias over time parallel visual working memory work, which shows evidence of a hierarchical organization of memory: items are more biased towards their center as uncertainty increases in order to increase the overall precision of retrieval (Brady & Alvarez, 2011; Lew & Vul, 2015; Orhan & Jacobs, 2013). Our results detected a similar gist-based bias in long-term memory consolidation. Moreover, in Experiment 2, after a long retention interval, the reported gist overweighted the outlier, whereas the item memories were biased towards the local gist which discounted the outlier. This finding also mirrors prior ensemble perception results that outliers are discounted or excluded in estimating summary statistics (De Gardelle & Summerfield, 2011) and suggests that the gist influencing item retrieval is not a simple average of the items. The results might reveal two different sampling strategies for gist extraction. Because participants had explicit knowledge about the outlier, they might have given more weight to the outlier in explicitly recalling and reporting the gist, similar to the change in the pattern by inconsistent items observed in long-term memory work (Richards et al., 2014). In contrast, the local center that influenced the items might reflect an implicit representation with a sampling strategy discounting the outlier, consistent with findings in perception work (De Gardelle & Summerfield, 2011; Haberman & Whitney, 2010). Our results suggest that visual working memory and long-term memory might be underpinned by a similar reconstructive mechanism and open up new directions to bridge the two fields.

Future research can be done to test the generality of our findings to other domains of human cognition. It would be interesting to explore whether our findings, which considered gist memory as a spatial average, would generalize to a broader definition of gist, such as gist-like memory for events (Moscovitch, Cabeza, Winocur, Nadel, 2016). For example, when first learning what a birthday party is from attending a few, the “gist” representation of a birthday party may be dominated by memory for a few parties, but over time the gist becomes a more stable representation that can influence retrieval of those specific birthday party events. In addition, the dissociation of gist-based bias in Experiment 2 also mirrors the dissociable implicit and explicit attitude in social categories (Gawronski & Bodenhausen, 2006). Future work could further disentangle these processes in long-term memory consolidation, which could enlighten our understanding of the cognitive mechanism underlying the formation of gist in social categories.

One limitation of the current experiments is that the testing order (i.e., gist memory before item memory) might have encouraged the retrieval of the items to be consistent with the gist (Tversky & Kahneman, 1974; Mutluturk & Boduroglu, 2014). We initially chose this order because we were most interested in the change in gist representation and wanted to minimize the influence of item memories on gist estimation in later recall. This order could have encouraged a positive relationship between item and gist memory, as retrieving the gist could have facilitated memory for items. However, because the order is the same for the three different delay intervals, the changes in item memory and bias across delay groups could not be simply a result of the order of testing. Because the correlation was derived from the same item memory responses used to generate the bias results, and we observed changes in bias over time but no change in the relationship between item memory and gist, it is unlikely that the correlation was completely independent of the increase in bias and was solely driven by the testing order. Nevertheless, future studies with randomized testing order will be helpful to control the influence of testing order.

In summary, we have shown that memory for individual items and memory for the gist of a set of items changed over the course of long-term memory consolidation. We propose that the gist that was initially extracted from item memories gradually started to guide item memory retrieval over longer durations as their relative memory strength changed. These findings bridge research in areas of cognitive science ranging from perception and working memory to episodic and semantic memory, providing important new insights into our ability both to learn about distinct events and to generalize across similar experiences.

## Methods

### Experiment 1

#### Participants

In Experiment 1, we recruited 147 members of the University of Pennsylvania community (18-30 years old; normal or corrected to normal vision) to participate in the experiment for monetary compensation. Participants selected to sign up for a second session that followed their first session by either 24 hours, one week, or one month^2^. Sample size was based on Experiment 2 which was conducted first. We excluded 10 participants because of low performance on Session 1 (i.e. reported gist was out of the scope of the learned landmarks) and then 7 participants because of individual and gist performance of any sessions lower than 3 *SD* below average. Our reported results thus include 130 participants, with 44 participants in the 24-hours group (age: *M* = 21.3, *SD* = 2.9, gender: 61% females), 43 participants in the one-week group (age: *M* = 21.9, *SD* = 2.6, gender: 67% females), and 43 participants in the one-month group (age: *M* = 21.4, *SD* = 2.0, gender: 74% females). All procedures were approved by University of Pennsylvania IRB.

##### Procedure

The experimental procedure is displayed in Fig. 1b. All participants completed Sessions 1 and 2; the only difference between groups was the time delay between sessions. Session 1 included training and testing. During training, participants were trained to retrieve six landmark locations consecutively on a laptop until their retrieval error for each landmark was fewer than 80 pixels in any direction. 80 pixels was chosen to be the criterion because it was less than 1/2 of the shortest distance between any pairs of the encoded locations, and thus would ensure that participants could differentiate the locations in recall. The training included three phases. In Phase 1, the landmarks appeared on the screen one at a time, and participants were required to click on each landmark to proceed. Fig 1A illustrates the landmark locations; note, on each trial, only one location was presented (never the full map), and the center of the encoded locations was never presented to participants. In Phase 2, we asked participants to recall the location for each landmark by clicking on the screen when given its name as a cue, and we gave them feedback about their guesses: participants had 3 attempts to recall each landmark location. For each attempt, if the distance between the recalled location and retrieved location satisfied the training criterion (i.e., 80 pixels), the correct location would be shown on the screen; otherwise a message would be prompted that their attempt was incorrect. The correct location would be shown on the screen after three incorrect attempts. In Phase 3, participants recalled each landmark consecutively without feedback, one at a time. If each of the retrieved landmarks fell in the range of 80 pixels, the participant could proceed to testing; if not, the participant was redirected back to Phase 2 to receive more training. After participants reached the training criterion, they completed ten unrelated arithmetic problems, in order to minimize potential influences from working memory. Finally, participants were tested on their memory of the locations: They indicated their guess about the center of the landmarks (gist memory test), with an instruction “Indicate the center (average location) of the landmarks you have seen”. Then, they separately recalled each landmark location (item memory test, which was identical to the recall procedure in Phase 3). In both tests, participants were incentivized to be accurate through bonus payments. They would receive a bonus of 1 dollar for the gist memory test if their error was within 100 pixels and a bonus of 1 dollar for the item memory test if their average error across all items was within 80 pixels. All trials were self-paced. The total time for Session 1 was approximately 12 minutes.

After 24 hours, one week, or one month, participants returned for Session 2. Session 2 was identical to the testing phase of Session 1, in which participants first reported the center and then the location of each landmark. Trials were again self-paced. The total time for Session 2 was approximately 5 minutes. Participants could choose to quit the experiment after Session 1 and received 10 dollars for their time, otherwise they would be paid after Session 2. The payment ranged from 16 to 20 dollars, depending on participants’ performance in their gist memory and item memory test.

### Error Measurement

In order to measure the accuracy for item memory (memory for each landmark), gist memory (reported memory for the center of the landmarks), and the estimated gist (center of all the retrieved items), we developed three error measurements as follows.

Item Memory Error (green line in Fig. 2a): The error for each item was defined as the Euclidean distance between the retrieved location for each landmark and its encoded location. Each participant’s item memory error was computed as the average error for the six landmarks. Chance performance is 348 pixels, which is determined by the average Euclidean distance between the center of the laptop screen and each encoded item location. Mathematically, this distance corresponded to the average error a participant would have if they guessed anywhere on screen.

Gist Memory Error (purple line in Fig. 2a): The error for gist memory was defined as the reported center error, which was the Euclidean distance between the participant’s reported center and the true center of all the encoded items. Chance performance is 216 pixels, which is the Euclidean distance between the center of the laptop screen and the true center of all encoded locations.

Estimated Gist Memory Error (blue line in Fig. 2a): The error for the estimated gist based on items was defined as the estimated center error, which was the Euclidean distance between the center of each participant’s retrieved item locations and the true center of all encoded locations. In other words, the estimated center can be thought of as what the participant’s gist estimate would be if it were directly computed by averaging across all retrieved item locations.

### Bias Measurement (Controlling for Error)

In order to measure the influence of gist on item memory, we developed a bias measurement as follows. Since the error analysis revealed a decrease in gist memory (i.e. reported center) after a month, there could be a difference in using reported center and using the true center of encoded items as bias center. We initially used the reported center as the center for the bias analysis. However, we also computed the bias index using the true center of encoded items to be consistent with common practices in ensemble perception research (Brady & Alvarez, 2011; Lew & Vul, 2015).

#### Absolute Bias

The bias towards the center for each retrieved item was defined as the relative difference in distance between a participant’s reported center and each landmark’s encoded location versus each landmark’s retrieved location. This relative difference was then divided by the error for that landmark: (Encoded Item - Reported Gist) - (Retrieved Item - Reported Gist) / (Encoded Item - Retrieved Item; Fig. 3b). Bias thus can range between −1 and 1 and bias > 0 indicates that item memory is biased towards the center while bias < 0 indicates that the item is biased away from the center. Each participant’s bias not controlling for error was computed as the average across the biases of the 6 landmarks.

#### Baseline Bias

When item memory error increases, the increased error will lead to a negative absolute bias value despite no meaningful bias away from the center of the landmarks (see Fig. 7 for an illustration).

**Fig. 7.**
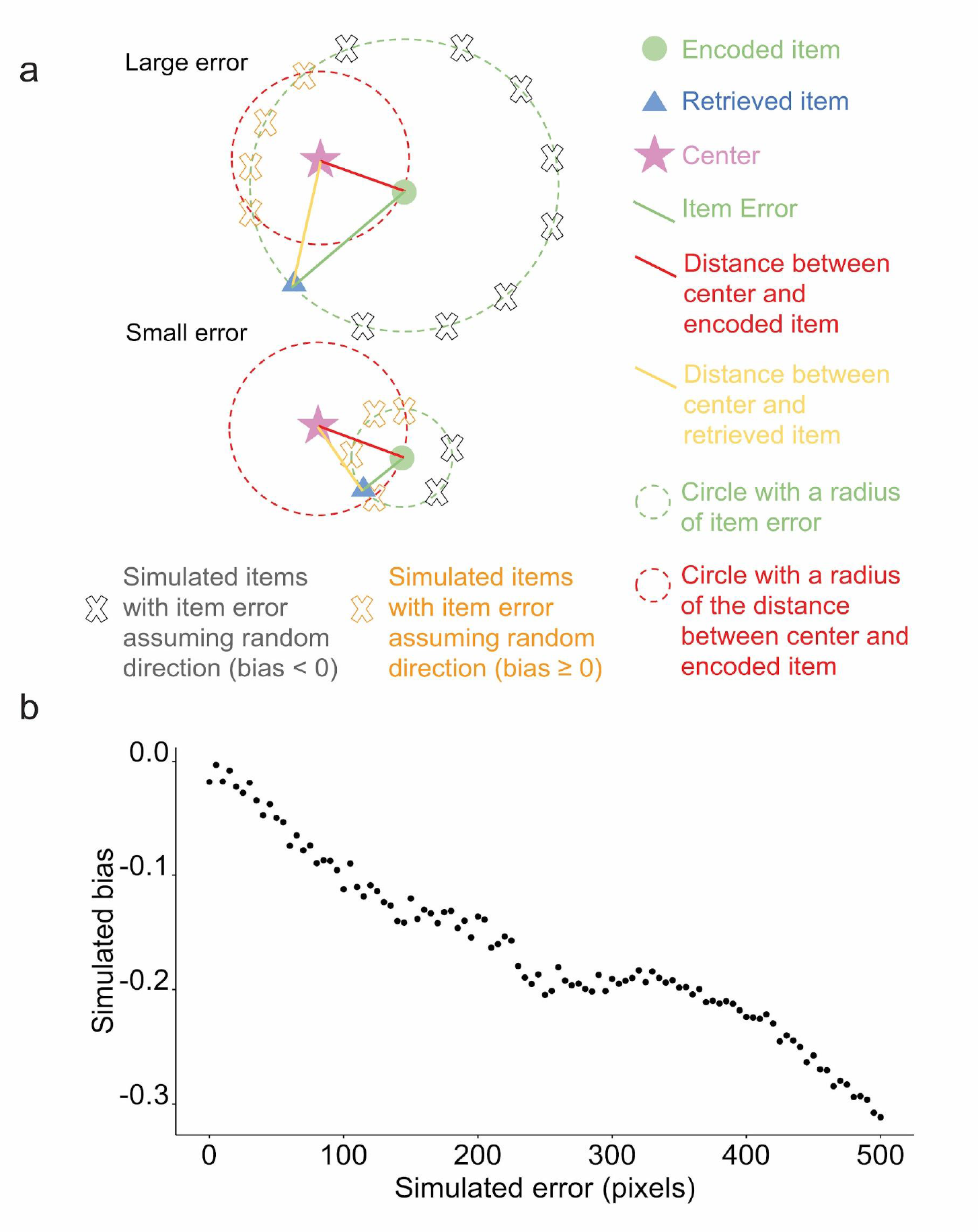
a) Items with large errors are more likely to have a negative absolute bias by chance, which is demonstrated in the figure in the following. First, the proportion of the arc with negative absolute bias in the circumference of simulated items is higher for items with large error. Because the distance between any points on the arc defined by the intersection between the green circle and red circle to the center will be shorter than the red distance, the points will all have an absolute bias value ≥ 0 (indicated by orange X marks), whereas the points outside of the arc will have a negative bias (indicated by grey X marks). Second, even though the retrieved item (blue triangle in the lower figure) with small error and the retrieved item with large error (blue triangle in the upper figure) are biased in exactly the same direction, the absolute bias for the retrieved item with the small error is positive whereas the other is negative, which demonstrates how a retrieved item with large error could cause negative bias without meaningfully being biased away from the center relative to its encoded location. b) A simulation based on the 6 encoded locations in Experiment 1 that shows that random error of retrievals not assuming direction was negatively correlated with bias.

Therefore, to control for the influence of error magnitude on our estimate of bias, we conducted a simulation to generate a baseline. We generated 1000 simulations for each participant. Each simulation consisted of six simulated retrieved items, corresponding to all the six landmark locations. For each location, we randomly generated a retrieved location based on the participant’s true error for this specific location, allowing angle to vary randomly across the simulations (Fig. 7b; gray cross). If a simulated location fell outside the boundaries of the screen, the algorithm generated a new location. The bias value for each of the 1000 simulations was the average across each simulation’s six retrieved locations. The baseline bias for each participant was the average across the 1000 simulations.

Bias = Absolute Bias - Baseline Bias: We subtracted the simulated bias from participants’ true bias, which resulted in a bias controlling for error. Bias > 0 indicates that item retrieval is biased towards the center, controlling for item memory error.

### Statistics

To examine whether gist memory persisted when memory for items decayed over time, we conducted a 3 (group: 24-hour, 1-week, and 1-month) X 2 (memory type: item, center) aligned ranks transformation ANOVA of the error change (Session 2 error values - Session 1 error values) and also two-tailed Mann-Whitney tests for between-group error change comparisons, because the data were not normally distributed as determined by a Shapiro-Wilk test. In order to examine whether there is a relation between item and gist memory, we used a linear model to evaluate the effects of estimated center error, delay group, and their interaction on reported center error.

In order to examine whether item retrievals were biased towards the center at any time points, we compared the bias value against 0 for each group at each session. In order to examine whether the bias controlling for error described above increased over time, we conducted a one-way ANOVA of change in bias (Session 2 bias values - Session 1 bias values) across the three groups. We used two-tailed t-tests for between-group comparisons. Reports were not multiple comparisons corrected.

### Swapped Items

In recalling the location for the landmarks, participants might “misbind” the label of a landmark and its location (e.g., indicate the location of the “restaurant” at the actual location for the “university” and vice versa). In order to test the potential influence of such errors on our results, we developed a criterion to identify pairs of items that were swapped, and we swapped them back to see if that changed the results. That is, for example, if (1) the retrieval for “restaurant” was closest to the encoded location for “university”, (2) the retrieval for “university” was closest to the encoded location for “restaurant”, (3) the retrievals were both within the range of both of the encoded locations (i.e. the distance between encoded “restaurant” and “university” / 2) and, (4) there were no other retrievals in this range, we then swapped the retrieved university and restaurant responses and used the swapped results for the analyses described above. We found that swapping the items did not change any of the reported results.

## Experiment 2

### Participants and procedure

We recruited 77 members of the University of Pennsylvania community (18-30 years old; normal or corrected to normal vision) to participate in the experiment for monetary compensation. Sample size was based on prior behavioral memory studies. We maximized data collection across two semesters. All procedures were approved by University of Pennsylvania IRB. All 77 participants received training and testing during Session 1 and reported item and center memories again after 24 hours (Session 2). Sessions 1 and 2 were identical with Experiment 1, except that in Experiment 2 during Session 1, participants were trained to retrieve eight landmark locations, one of which was a spatial outlier (see Fig. 4a), until their retrieval error for each landmark was fewer than 50 pixels (again, a distance less than 1/2 of the shortest distance between the pairs of encoded locations) in any direction. In addition, in Session 1 of Experiment 2, to streamline the session, item memory was derived from the item memory test in the last round of evaluation during training (Phase 3), which was immediately followed by the gist memory test (Fig. 4b). The time for Session 1 was approximately 25 minutes, which was longer than that for Experiment 1 because in Experiment 2, participants learned more locations and the training criterion was harder (50 pixels, as opposed to 80 pixels in Experiment 1).

After 32 to 57 days, 50 participants returned for Session 3 by email invitation. Session 3 was identical to Session 2 (i.e., participants reported their memory for the center and then each item). The time for Session 2 and 3 was approximately 10 minutes. Of the 50 participants who returned for the third session, 1 participant was excluded because their individual and gist performance for at least one session was lower than 3 *SD* below average. We did not exclude participants whose reported gist memory error was larger than the distance between the screen center and the true center at Session 1, as in Experiment 1, because in Experiment 2, a large gist error could be a meaningful result that reflects the overweighting of the outlier in reporting gist. We excluded 6 participants who placed the outlier where the majority of items were, which means the error of the outlier was larger than 573 pixels (i.e. the distance between the center of screen and outlier encoded location). The reason we excluded these participants was that in Experiment 2, if participants swapped the outlier with one of the other items, or simply put the outlier among the other items, this outlier swap would strongly inflate the bias value towards the global reported center, which does not necessarily reflect a true bias towards the center. Our reported results thus include 43 participants (age: *M* = 21.5, *SD* = 2.2, gender: 75% female)

### Error measurement

All error measures were calculated as in Experiment 1, except that the chance performance for individual items was 386 pixels (determined by the average Euclidean distance between the center of the laptop screen and each encoded item location) and the chance performance for gist memory was 223 pixels (determined by the average Euclidean distance between the center of the laptop screen and the true center of all encoded item locations).

### Bias Measurement (Controlling for Error)

We calculated bias as in Experiment 1, except that we additionally computed a local gist bias, which was a bias index using the local center (i.e. the center of the seven encoded locations excluding the outlier) as the bias center.

### Outlier weight estimation

In order to estimate the weight of the outlier on the reported gist memory, we developed a weight model as follows: For each participant at each session, a series of weights were applied to the encoded outlier location. The range of the weight of the outlier was from 0 to 1, with a stepwise increment of 0.0125. The weight for each of the other encoded items was assumed to be the same and would thus be (1 - outlier weight)/number of items that were not the outlier, ranging from 0 to 0.125. Based on these weights, 81 simulated centers were computed: when the outlier weight was 0, the simulated center would be a perfect local center ignoring the outlier; when the outlier weight was 0.125, the simulated center would be a perfect global center of all items, since there were 8 items in total; when the outlier weight was 1, the simulated center would be the outlier itself.

For each participant at each session, the Euclidean distance between each simulated center and reported center was computed, resulting in 81 distances. We used the weight that resulted in the smallest distance as the estimated weight of the outlier for that participant at that session.

### Statistics

As in Experiment 1, we conducted a 2 (delay: short retention of 1 day or long retention of 1-2 months) x 2 (memory type: item, center) aligned ranks transformation ANOVA with repeated measures of error change (short, defined by Session 2 error values - Session 1 error values, or long, defined by Session 3 error values - Session 1 error values) and pairwise Wilcoxon signed rank tests for error comparisons, since change in error was not normal as determined by a Shapiro-Wilk test.

As in Experiment 1, in order to examine whether there was a relation between item and gist memory, we used a linear mixed effects model on reported center error with fixed effects of delay (24 hours and 1 to 2 months), estimated center error, and their interaction, as well as a random effect of participant.

In order to examine whether the outlier was weighted more, the same, or less compared to other items, we compared the outlier weight values against the weight assuming all items to be equal (i.e. ⅛ = 0.125) with t-tests. In order to examine whether the weight of the outlier in gist memory changed over time, we used a paired t-test comparing the outlier weight change after a short retention interval (Session 2 outlier weight values - Session 1 outlier weight values) against the outlier weight change after a long retention interval (Session 3 outlier weight values - Session 1 outlier weight values).

As in Experiment 1, we compared the bias values controlling for error against 0 at each session by t-tests to examine whether item retrievals were significantly biased towards the reported center. In order to examine whether the item retrievals were increasingly biased towards the reported center over time, we conducted a paired t-test comparing the bias change after a short retention interval (Session 2 bias values - Session 1 bias values) against the bias change after a long retention interval (Session 3 bias values - Session 1 bias values). We conducted the same statistical analyses using the local center as a bias center. Reports were not multiple comparisons corrected.

## Acknowledgement

The authors gratefully acknowledge the data collected by Siqi Lin, Jennifer Nazario, Emily Potter and the helpful discussions with colleagues in Penn Psychology.

## Data availability

All data will be available at https://osf.io/jxme8/?view_only=049dcb1efaf44c3098040ba027f88115 upon the acceptance of this manuscript.

## Code availability

All scripts used to analyze the data will be available at https://osf.io/jxme8/?view_only=049dcb1efaf44c3098040ba027f88115 upon the acceptance of this manuscript.

---

**Supplementary Figure 1.**
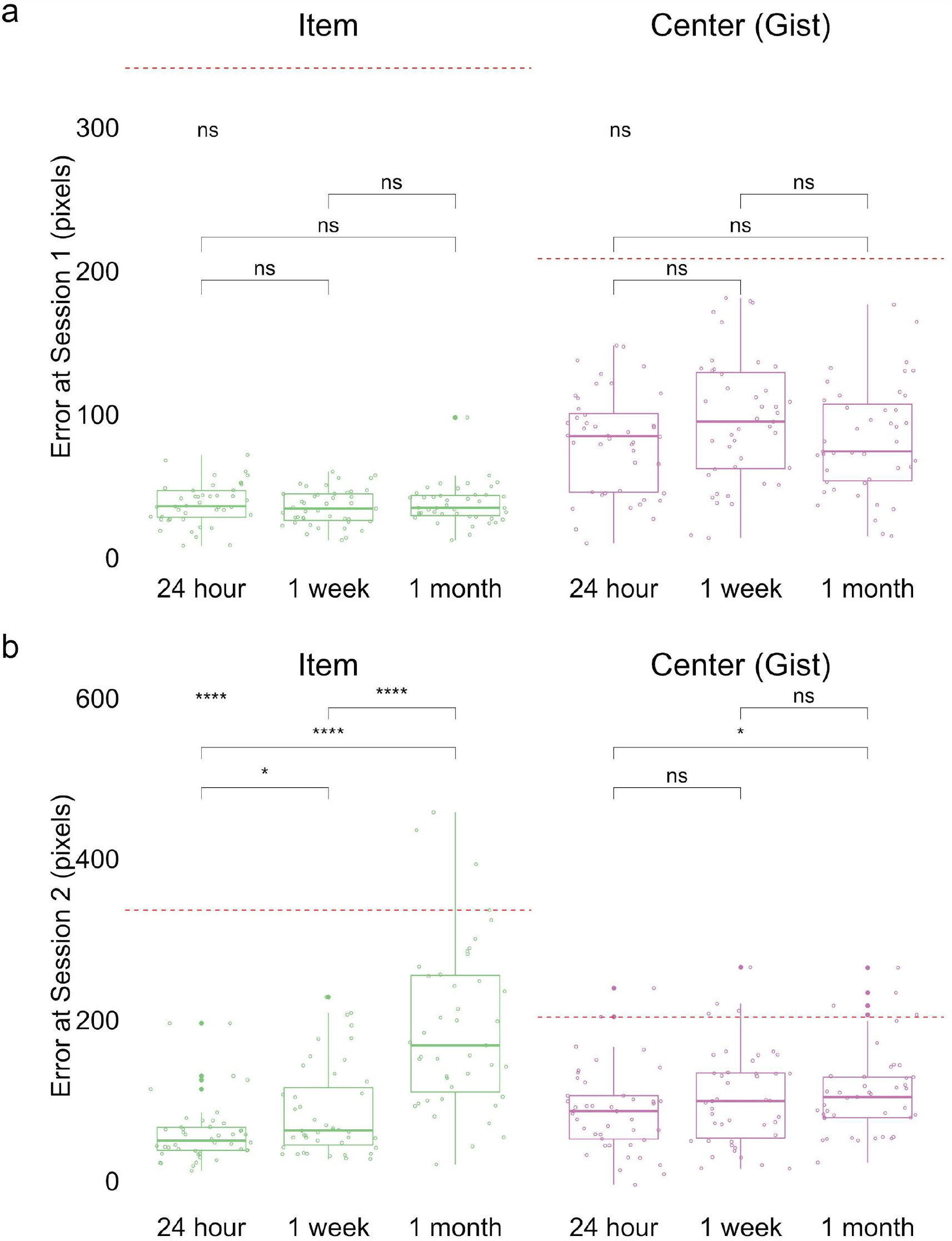
Experiment 1 error in item and gist (center) memory at Session 1 (a) and at Session 2 (b). * indicates *p <* .05, **** indicates *p* < .0001, and ns indicates p > .05 by t-tests between groups and ANOVA (top left). Red dashed lines indicate chance performance. The band indicates the median, the box indicates the first and third quartiles, the whiskers indicate ± 1.5 × interquartile range, and the solid points indicate outliers.

**Supplementary Figure 2.**
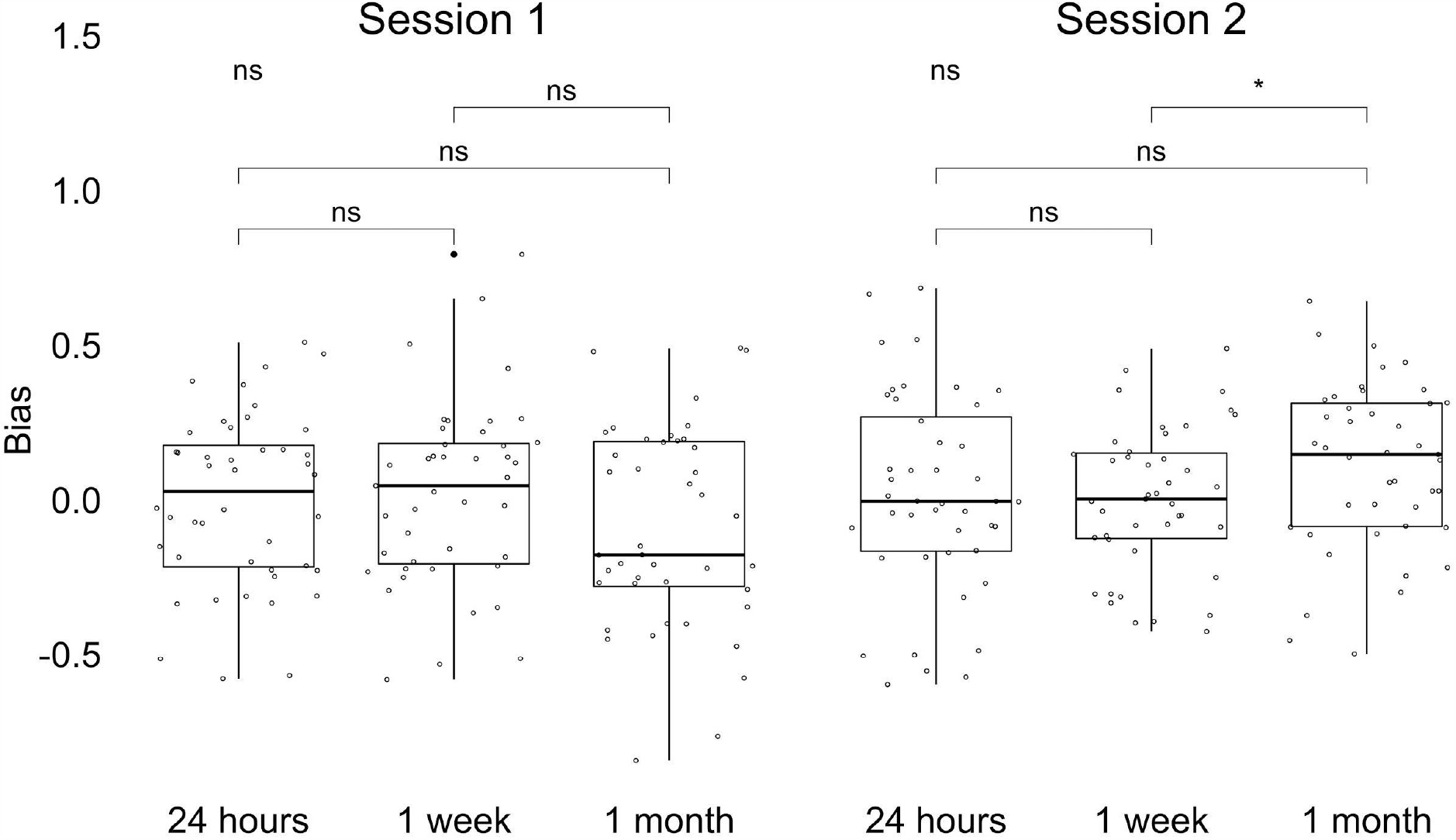
Experiment 1 Gist-based bias for three delay groups at Session 1 and Session 2. * indicates p < .05 of difference between groups by t-tests or ANOVA (top left). The band indicates the median, the box indicates the first and third quartiles, the whiskers indicate ± 1.5 × interquartile range, and the solid points indicate outliers.

**Supplementary Figure 3.**
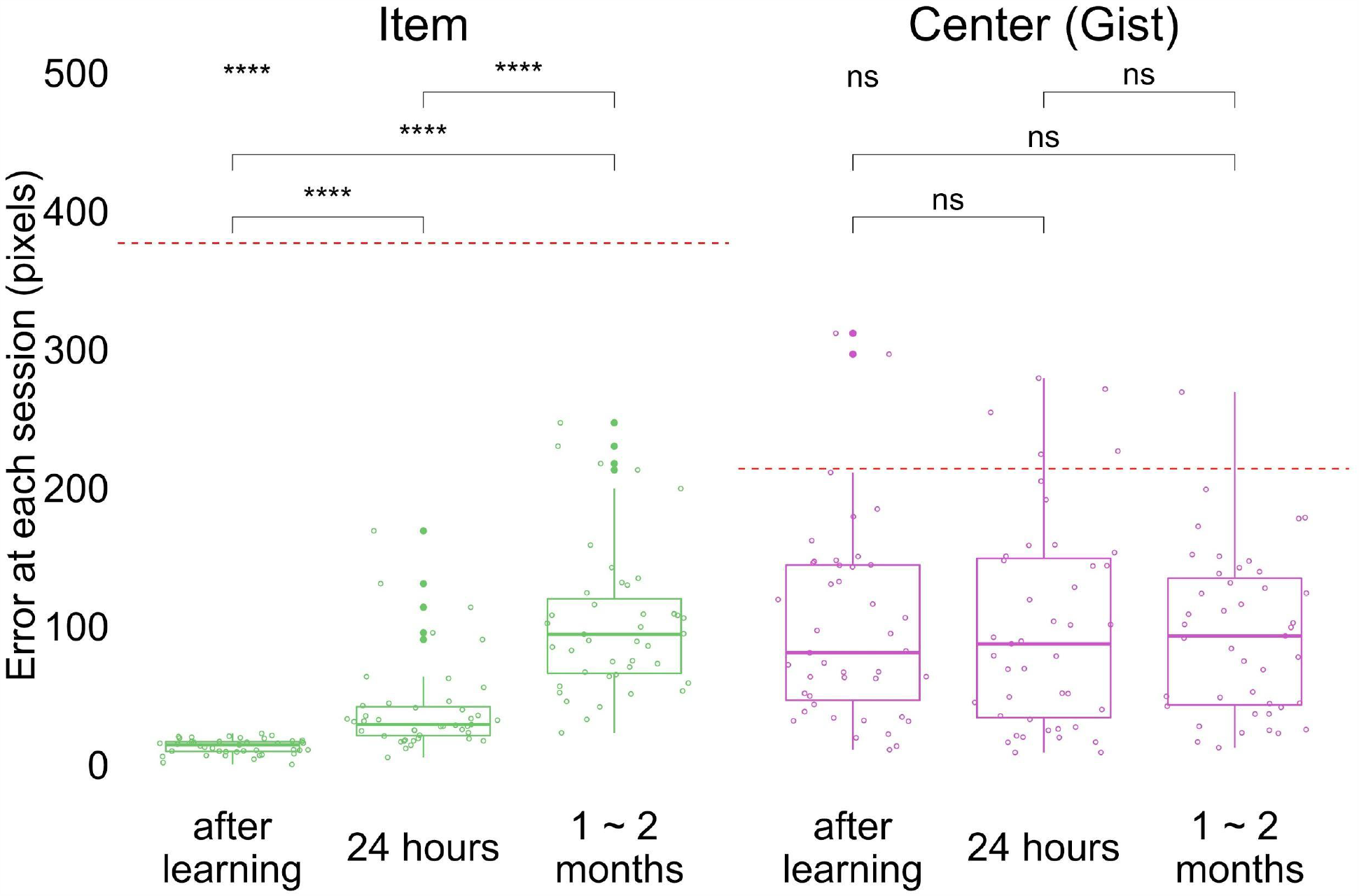
Experiment 2 error in item and gist (center) memory at each session. Red dashed lines indicate chance performance. **** indicates p < .0001 by paired t-tests and ANOVA (top left). The band indicates the median, the box indicates the first and third quartiles, the whiskers indicate ± 1.5 × interquartile range, and the solid points indicate outliers.

**Supplementary Figure 4.**
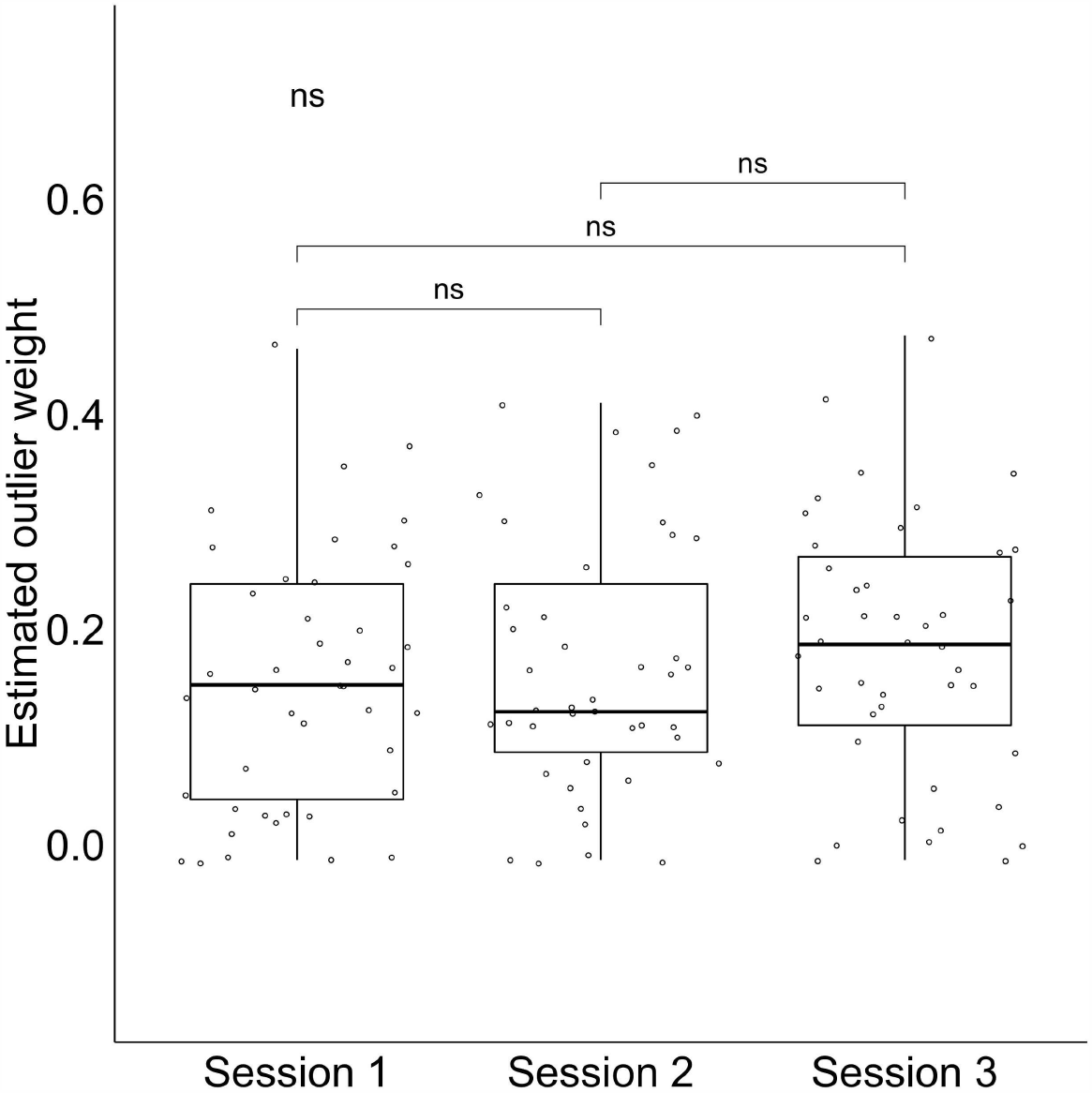
Experiment 2 outlier weight values at each session. The band indicates the median, the box indicates the first and third quartiles, the whiskers indicate ± 1.5 × interquartile range.

**Supplementary Figure 5.**
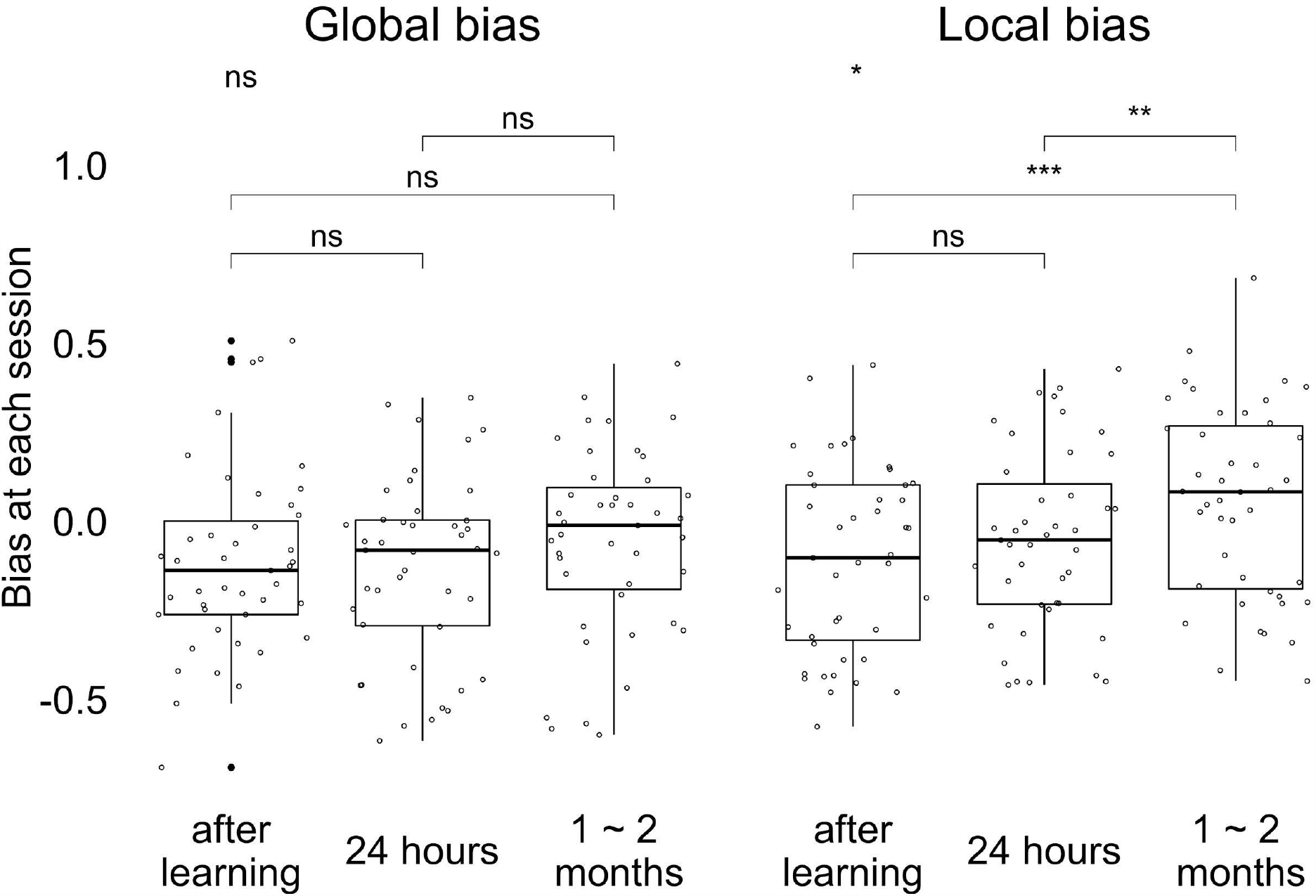
Experiment 2 Gist-based biases at each session. * indicates p < .05, ** indicates *p* < .01, and *** indicates *p* < .001 by paired t-tests and ANOVA (top left). The band indicates the median, the box indicates the first and third quartiles, the whiskers indicate ± 1.5 × interquartile range, and the solid points indicate outliers.

1 To be consistent with common practices in ensemble perception research (Brady & Alvarez, 2011; Lew & Vul, 2015), we repeated the bias analysis using the true center of encoded items (big green circle in Fig 2a) instead of the reported center, and we found the same pattern, with a main effect of group, *F*(2, 127) = 13.26, *p* < .001, a greater change in bias between the 24-hour and 1-month groups, *t*(71.88) = 3.27, *p* = 0.001, and also between the 1-week and 1-month groups *t*(75.18)= 3.79, *p* < 0.001, but not between the 24-hour and 1-week groups, *t*(84.21) = −0.78, *p* = .44. Again, only the bias at 1 month was significantly greater than 0, t(42) = 3.94, p < .001. Items memory also became more biased towards the true center of encoded locations after one month.

2 Because participants were not randomly assigned into three different delay conditions, a difference in expectation may influence their learning and consolidation. We did not find evidence, however, for any differences in behavior between groups at initial learning (Supplementary Fig. 1). Also, the results from Experiment 1 replicated in Experiment 2 with a within-subject design.

## Reference

Alvarez, G. A. (2011). Representing multiple objects as an ensemble enhances visual cognition. Trends in Cognitive Sciences, 15(3), 122–131. https://doi.org/10.1016/j.tics.2011.01.003

Ariely, D. (2001). Seeing sets: representation by statistical properties. Psychological Science, 12(2), 157–162.

Ariely, D., & Carmon, Z. (2003). Summary assessment of experiences: the whole Is different from the sum of its parts. Retrieved from https://www.ebsco.com/terms-of-use

Brady, T. F., & Alvarez, G. A. (2011). Hierarchical encoding in Visual Working Memory: Ensemble Statistics Bias Memory for Individual Items. Psychological Science, 22(3), 384– 392.

Brady, T., Schacter, D., & Alvarez, G. (2015). The adaptive nature of false memories is revealed by gist-based distortion of true memories. Preprint, 15(12), 948. https://doi.org/10.1167/15.12.948

De Gardelle, V., & Summerfield, C. (2011). Robust averaging during perceptual judgment. Proceedings of the National Academy of Sciences of the United States of America, 108(32), 13341–13346. https://doi.org/10.1073/pnas.1104517108

Gawronski, B., & Bodenhausen, G. V. (2006). Associative and propositional processes in evaluation: an integrative review of implicit and explicit attitude change. Psychological Bulletin, 132(5), 692–731. https://doi.org/10.1037/0033-2909.132.5.692

Haberman, J., & Whitney, D. (2010). The visual system discounts emotional deviants when extracting average expression. Attention, Perception & Psychophysics, 72(7), 1825–1838. https://doi.org/10.3758/APP.72.7.1825.The

Hemmer, P., & Steyvers, M. (2009). A Bayesian account of reconstructive memory. Topics in Cognitive Science, 1(1), 189–202. https://doi.org/10.1111/j.1756-8765.2008.01010.x

Huttenlocher, J., Hedges, L. V., & Duncan, S. (1991). Categories and particulars: prototype effects in estimating spatial location. Psychological Review, 98(3), 352–376. https://doi.org/10.1037/0033-295X.98.3.352

Huttenlocher, J., Hedges, L. V., & Vevea, J. L. (2000). Why do categories affect stimulus judgment? Journal of Experimental Psychology: General, 129(2), 220–241. https://doi.org/10.1037/0096-3445.129.2.220

Lau, H., Tucker, M. A., & Fishbein, W. (2010). Daytime napping: effects on human direct associative and relational memory. Neurobiology of Learning and Memory, 93(4), 554–560. https://doi.org/10.1016/j.nlm.2010.02.003

Lew, T. F., & Vul, E. (2015). Ensemble clustering in visual working memory biases location memories and reduces the Weber noise of relative positions. Journal of Vision, 15(4), 1–14. https://doi.org/10.1167/15.4.10

Lutz, N. D., Diekelmann, S., Hinse-Stern, P., Born, J., & Rauss, K. (2017). Sleep supports the slow abstraction of gist from visual perceptual memories. Scientific Reports, 7(1), 1–9. https://doi.org/10.1038/srep42950

Moscovitch, M., Cabeza, R., Gordon, W., & Nadel, L. (2016). Episodic memory and beyond: the hippocampus and neocortex in transformation. Annual Review of Psychology, 67, 105–134. https://doi.org/10.1016/j.physbeh.2017.03.040

Mutluturk, A., & Boduroglu, A. (2014). Effects of spatial configurations on the resolution of spatial working memory. Attention, Perception, and Psychophysics, 76(8), 2276–2285. https://doi.org/10.3758/s13414-014-0713-4

Orhan, A. E., & Jacobs, R. A. (2013). A probabilistic clustering theory of the organization of visual short-term memory. Psychological Review, 120(2), 297–328. https://doi.org/10.1037/a0031541

Posner, M. I., & Keele, S. W. (1970). Retention of abstract ideas. Journal of Experimental Psychology, 83(2p1), 304–308. https://doi.org/10.1037/h0028558

Richards, B. A., Xia, F., Santoro, A., Husse, J., Woodin, M. A., Josselyn, S. A., & Frankland, P. W. (2014). Patterns across multiple memories are identified over time. Nature Neuroscience, 17(7), 981–986. https://doi.org/10.1038/nn.3736

Richter, F. R., Bays, P. M., Jeyarathnarajah, P., & Simons, J. S. (2019). Flexible updating of dynamic knowledge structures. Scientific Reports, 9(1), 1–15. https://doi.org/10.1038/s41598-019-39468-9

Schacter, D. L., Guerin, S. A., & St. Jacques, P.L. (2011). Memory distortion: An adaptive perspective. Trends in Cognitive Sciences, 15(10), 467–474. https://doi.org/10.1016/j.tics.2011.08.004

Sekeres, M. J., Winocur, G., & Moscovitch, M. (2018). The hippocampus and related neocortical structures in memory transformation. Neuroscience Letters, 680(May), 39–53. https://doi.org/10.1016/j.neulet.2018.05.006

Squire, L. R., Genzel, L., Wixted, J. T., & Morris, R. G. (2015). Memory consolidation. Cold Spring Harbor Perspectives in Biology, 7(8), a021766.

Sweegers, C. C. G., & Talamini, L. M. (2014). Generalization from episodic memories across time: A route for semantic knowledge acquisition. Cortex, 59(49–61). https://doi.org/10.1016/j.cortex.2014.07.006

Tompary, A., & Thompson-schill, S. L. (2019). Semantic influences on episodic memory distortions. Proceedings of the 41st Annual Conference of the Cognitive Science Society, 1124–1130.

Tompary, A., Zhou, W., & Davachi, L. (2020). Schematic memories develop quickly, but are not expressed unless necessary. Scientific Reports, 1–17. https://doi.org/10.31234/osf.io/k4fea

Tversky, A., & Kahneman, D. (2019). Judgment under uncertainty. Foundations of Cognitive Psychology, 185. https://doi.org/10.7551/mitpress/3080.003.0038

Whitney, D., & Yamanashi Leib, A. (2018). Ensemble perception. Annual Review of Psychology, 69(1), 105–129.

Winocur, G., & Moscovitch, M. (2011). Memory transformation and systems consolidation. Journal of the International Neuropsychological Society, 17(5), 766–780. https://doi.org/10.1017/S1355617711000683

